# Controlling minimal and maximal hook-length of the bacterial flagellum

**DOI:** 10.1101/2020.03.25.007062

**Authors:** Alina Guse, Manfred Rohde, Marc Erhardt

**Author notes:** Correspondence to: Marc Erhardt, **Email:**.

## Abstract

Hook-length control is a central checkpoint during assembly of the bacterial flagellum. During hook growth, a 405 amino acids (aa) protein, FliK, is intermittently secreted and thought to function as a molecular measuring tape that, in *Salmonella*, controls hook-length to 55 nm ± 6 nm. The underlying mechanism involves interactions of both the α-helical, N-terminal domain of FliK (FliK_N_) with the hook and hook cap, and of its C-terminal domain with a component of the export apparatus. However, various deletion mutants of FliK_N_ display uncontrolled hook-length, which is not consistent with a ruler mechanism. Here, we carried out an extensive deletion analysis of FliK_N_ to investigate its contribution in the hook-length control mechanism. We identified FliK_N_ mutants deleted for up to 80 aa that retained wildtype motility. However, the short FliK variants did not produce shorter hook-lengths as expected from a physical ruler. Rather, the minimal length of the hook depends on the level of hook protein production and secretion. Our results thus support a model in which FliK functions as a hook growth terminator protein that limits the maximal length of the hook, and not as a molecular ruler that physically measures hook-length.

## Introduction

Many bacteria use a complex nanomachine, called the flagellum, to propel themselves through liquid environments. Flagella consist of three main structures, (i) a membrane-spanning basal body complex, which is connected by (ii) a flexible hook to (iii) the long rigid filament (1). The hook adapts a curved structure composed of approximately 120 subunits of the same protein, FlgE, reaching a length of 55 nm (± 6 nm) in *Salmonella enterica* (2–4). We recently showed that the physiological hook-length of 55 nm is optimal for the formation of the flagellar filament bundle in *Salmonella*, as too short hooks might be too stiff and too long ones might buckle because of a too high flexibility, thereby disrupting the filament bundle also in the absence of chemotactic stimuli (5).

Hook-length control has been intensely studied over the years as unraveling the precise underlying mechanism is complex. Even more so, considering that the control of hook-length is directly connected with substrate specificity switching of the entire flagellar protein secretion machinery (6). A type-III secretion system (T3SS) secretes most building blocks of the flagellum. The primary export gate is made of FliP, FliQ and FliR, which form an unique helical pore complex; and FlhA and FlhB, which make up the substrate docking platform and are involved in energy transduction (7–11). Substrate specificity switching of the T3SS describes the ability of this machinery to switch from recognizing early substrates, such as the rod and hook subunits, to accepting and secreting late substrates, including the hook-filament junction proteins, as well as the filament subunits (6, 9). This enables the cell to coordinate gene expression with the spatiotemporal assembly of sub-parts of the flagellum. The time when the substrate specificity switch occurs has been shown to be primarily dependent on two proteins, a protein controlling hook-length, FliK, and the T3SS component FlhB (3, 4, 6). Interactions between the C-terminal domains of both proteins is proposed to induce conformational rearrangements in FlhB and FlhA resulting in the switch from early to late substrate secretion (12, 13). Therefore, controlling hook-length by FliK and subsequent interaction with FlhB constitutes an important checkpoint during assembly of the flagellum. The α-helical, N-terminal domain of FliK (FliK_N_) harbors the peptide secretion signal within the first 40 aa and is thought to be responsible for hook-length measurement (5, 14, 15). A homologous protein (YscP in *Yersinia*, InvJ in *Salmonella*) controls needle-length in the closely related injectisome system (16, 17). Deletions or insertions in the N-terminal domain of the proteins responsible for hook-length control in the flagellum and needle-length control in the injectisome resulted in shorter or longer hooks and needles, respectively (3, 5, 15, 17, 18). This led to the model that FliK and homologs act as molecular ruler proteins, which physically measure hook/needle length. However, several deletions of the N-terminal domain of FliK (FliK_N_) were previously shown to result in the formation of extremely long hooks of uncontrolled length, called polyhooks (19). To our knowledge, no functional deletions in FliK producing a protein shorter than 363 aa have been described so far (5, 15). Here, we show that deletions of up to 80 aa in FliK_N_ (FliK length 325 aa) retain wildtype motility and result in hooks of controlled length. However, independent from the length of FliK_N_, the hook grows to a minimal hook-length of 45 nm. Hook-length control in the short FliK_N_ mutants can be achieved by slowing down hook polymerization. We conclude that FliK controls the maximal, but not minimal length of the flagellar hook and thus functions as a terminator of hook growth to limit its physiologically relevant length.

## Results

### Stepwise deletion of the N-terminal domain of FliK

FliK is a 405 aa protein and consists of a rather unstructured N-terminal domain (aa1-180, FliK_N_) that includes its peptide secretion signal (aa1-40), and a compactly folded C-terminal domain (aa206-405) connected by a flexible linker (aa181-205) (14). We first deleted 20, 40, 60, 80, 100, 120 and 140 aa regions within aa41-180 of FliK_N_. This resulted in various FliK proteins ranging from 385 to 265 aa in length. Most deletions had no effect on motility compared to the wildtype (WT), while some slightly reduced motility to about 85% of the WT (Fig. 1A+B and Fig. S1). We tried to further shorten FliK by deleting part of the region that is thought to harbor the secretion signal (Δaa21-40, Δaa31-40). As expected, these deletions resulted in a severe motility defect and were not included in further analysis (Fig. S2).

**Figure 1.**
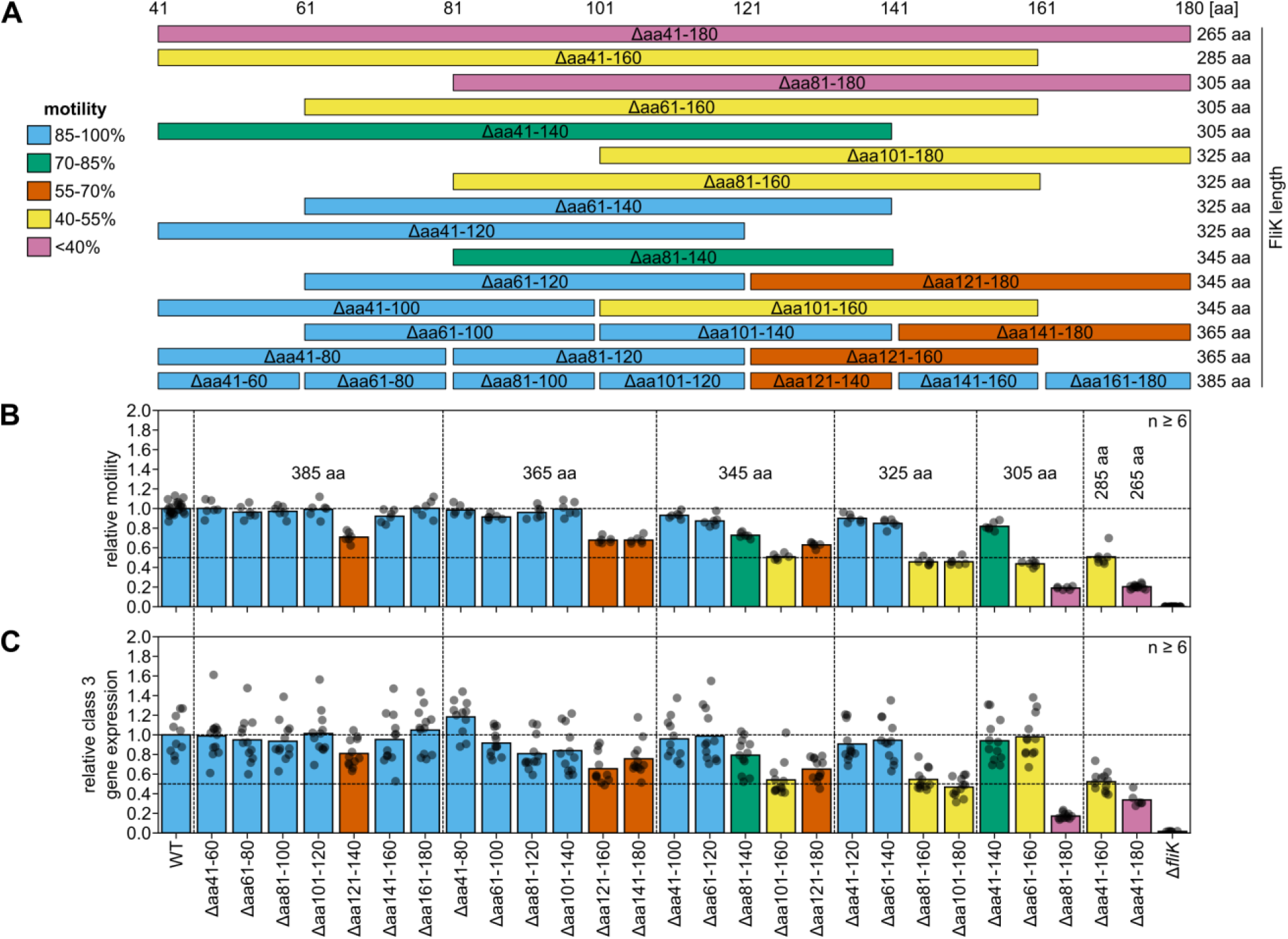
Motility performance and substrate specificity switching capability of FliK_N_ deletion variants. (A) Schematic overview of FliK_N_ deletion variants. Horizontal bars represent different deletions within the N-terminal domain of FliK (aa41-180). Colors correspond to the motility performance relative to the WT in percent as quantified in panel B. (B) Motility phenotype of the WT (FliK_405_) and FliK_N_ deletion variants analyzed using soft-agar swim plates. Relative motility represents the motility swarm diameters normalized to the WT strain. Representative motility plates are shown in Supporting Information Fig. S1. (C) Flagellar class 3 gene expression as a reporter of the substrate specificity switch capability in the WT (FliK_405_) and FliK_N_ deletion variants. Flagellar biosynthesis was synchronized using an anhydrotetracycline-inducible flagellar master regulator (P*_tet_*-*flhDC*) and analyzed using a P*_motA_*-*luxCDABE* reporter. The relative class 3 gene expression was determined by measuring luminescence 105 min after induction of FlhDC expression and is reported normalized to the WT strain. The bar graphs represent the mean of n≥6 biologically independent replicates. Replicates are shown as individual data points. WT, wildtype FliK_405 aa_; RLU, relative light units; aa, amino acids.

Most crucial for the motility function were deletions spanning the region between aa121-160, resulting in 43% to 70% motility compared to the WT. Interestingly, a deletion within this region (Δaa141-160) displayed 90% of WT motility. A similar deletion mutant in the region aa121-160 was investigated in a previous study (Δaa129-159) and displayed a less efficient interaction between FliK and the hook subunit FlgE, resulting in the formation of polyhooks with attached filaments (20). Deleting the regions between aa121-140 and aa81-140 reduced motility to 71% and 73%, respectively, whereas a deletion of aa61-140 and aa101-140 had only a mild (85% of WT) or no effect (100% of WT).

Thus, we concluded that both the length and position of the deletion within FliK_N_ were important for FliK function. We next investigated the timing of the substrate specificity switch. For this, we synchronized the onset of flagella biosynthesis using inducible expression of the flagellar master regulator FlhDC (P*_tetA_-flhDC*) and measured expression of flagellar genes from a class 3 promoter using a previously described reporter system (P*_motA_*-*luxCDABE*) (21–23). Expression from class 3 promoters occurs only upon termination of hook growth and the subsequent substrate specificity switch, which results in secretion of the anti-σ factor FlgM. Secretion of FlgM releases the flagellum-specific σ-factor σ^28^, which activates gene expression from class 3 promoters (24). Measuring gene expression from P*_motA_* after synchronization of flagella synthesis therefore allows to estimate the timing of the substrate specificity switch. We found that FliK_N_ deletion mutants with a strong motility defect exhibited a delay in the substrate specificity switch (Fig. 1C).

We next characterized the motility, flagellation and hook-length phenotype of deletion mutants starting from aa41 (Δaa41-60, Δaa41-80, Δaa41-100, Δaa41-120, Δaa41-140, Δaa41-160, Δaa41-180) in more detail. Hereafter, we refer to the individual deletions by the length of the resulting FliK molecule (i.e. FliK_385_ to FliK_265_). Motility of FliK_365_ to FliK_325_ was slightly reduced (90-97% of the WT) (Fig. 2A and Fig. S3A). The motility of FliK_305_, FliK_285_ and FliK_265_ was decreased to 83%, 51% and 20% of the WT, respectively. We next determined whether the number of flagella per cell is accountable for the observed slight motility defect of FliK_305_ (Fig. 2B and Fig. S3B).

**Figure 2.**
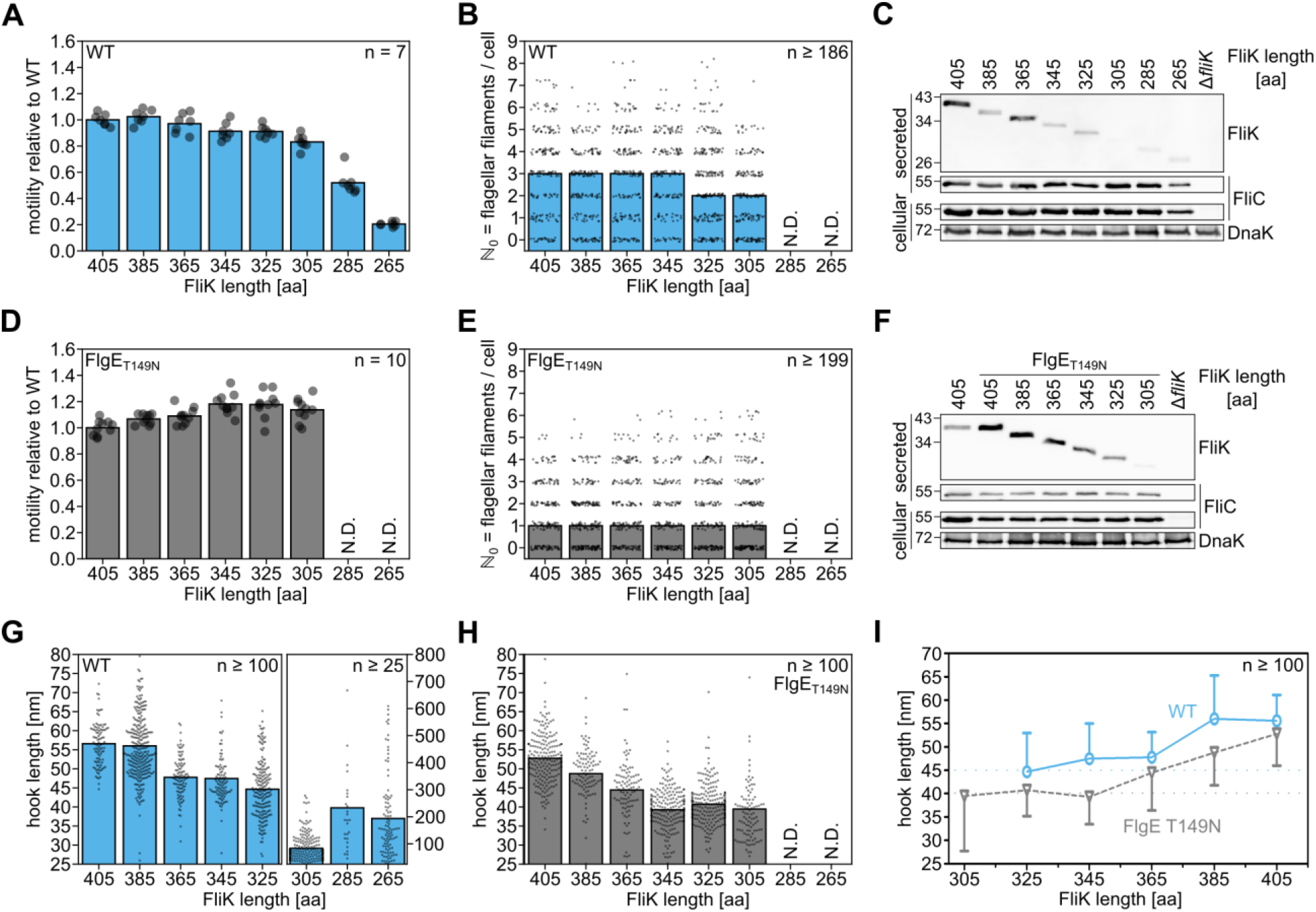
Effect of FliK_N_ deletion variants on motility, flagellation and hook-length control. Motility performance, flagellation and flagellar protein secretion of FliK variants were analyzed in the WT (A-C) and in the slow-hook polymerization mutant (FlgE_T149N_) background at 37 °C or 30 °C, respectively (D-F). (A+D) Motility phenotype of FliK_N_ deletion variants in the WT or FlgE_T149N_ background analyzed using soft-agar swim plates. Relative motility represents the motility swarm diameters normalized to the respective WT-FliK_405 aa_ strain. The bar graphs represent the mean of n=7 (WT) or n=10 (FlgE_T149N_) biologically independent replicates per strain. Replicates are shown as individual data points. (B+E) Flagellation of FliK_N_ deletion variants in the WT or FlgE_T149N_ background analyzed using anti-flagellin immunostaining. The bar graphs represent the median flagella per cell of n≥186 (WT) or n≥199 (FlgE_T149N_) individual bacteria analyzed per strain. The number of flagella per cell ℕ_0_ are shown as individual data points for each strain. (C+F) Cytoplasmic and secreted protein levels of the molecular ruler protein FliK and flagellin FliC of FliK_N_ deletion variants in the WT or FlgE_T149N_ background. DnaK served as loading and cell lysis control. The size of a protein ladder in kDa is indicated on the left. (G+H) Mean hook-length of FliK_N_ deletion variants in the WT (G) and FlgE_T149N_ (H) background. Hook-lengths of n≥25 isolated hook-basal body complexes were determined. Hook-length distributions for the individual strains are shown in Supporting Information Fig. S4 and S6. (I) Mean hook-length ± SD of FliK_N_ deletion variants in the WT [blue, solid line] or FlgE_T149N_ [grey, dashed line] background. Hook-lengths of n≥100 isolated hook-basal body complexes were determined. WT, wildtype; aa, amino acids; N.D., not determined.

FliK_305_ displayed a slight decrease in flagella per cell compared to the other mutants and the WT. Secretion analysis showed that all FliK variants were secreted into the culture supernatant, although the short variants FliK_305_, FliK_285_ and FliK_265_ appeared to be secreted less (Fig. 2C), which might be attributed to a decreased detection efficiency of our polyclonal α-FliK antibodies. However, we also cannot rule out decreased stability of the FliK deletion mutants as suggested before for other FliK variants (15, 18). We next purified hook-basal body (HBB) complexes of selected FliK mutants to determine their hook-lengths and analyze the ability to control hook-length (Fig. 2G+I and Fig. S4). FliK_385_ to FliK_325_ displayed hooks of controlled length, whereas FliK_305_ and shorter exhibited a broad hook-length distribution, even though FliK_305_ only exhibited a mild motility defect. Surprisingly, decreasing FliK length did not result in correspondingly shorter hook-lengths as would be expected from a mechanism where FliK determines hook-length as a physical molecular ruler. We found that below a certain FliK length (FliK_325_), hook-length did not decrease below a minimal length of 45 nm. We therefore concluded that another mechanism contributes to hook-length control.

### Slow hook polymerization decreases hook-length in short FliK variants

It has previously been shown that the ratio of hook subunit FlgE production compared to FliK production is important for proper hook-length control. Shorter hooks have been observed under conditions when FliK was overproduced (25, 26), FlgE was underproduced (27) or in a hook polymerization defective mutant (2). In addition, longer hooks have been observed under conditions when FlgE was overproduced or FliK underproduced (25, 27). Accordingly, we speculated that the level of FlgE production, secretion and polymerization into the growing hook contributes to the observed minimal hook-length of 45 nm. We therefore manipulated the rate of hook growth using the slow-hook polymerization variant FlgE_T149N_ and investigated its effect on hook-length control in our FliK_N_ deletion variants. The FlgE_T149N_ mutation was originally isolated as a temperature-sensitive mutant which produced a population of shorter hooks at elevated temperature, but also hooks of WT length at 30 °C (2, 28, 29). As previously shown, the FlgE_T149N_ mutant displayed a motility of 50% compared to the WT at 30 °C (2). The FliK_N_ variants investigated here displayed no difference in motility compared to the parental FlgE_T149N_ mutant at 30 °C (Fig. 2D and Fig S5A). Consistently, all short FliK_N_ FlgE_T149N_ variants produced a similar number of flagella per cell as analyzed by flagella immunostaining (Fig. 2E and Fig S5B), but overall less flagella than in the WT background (Fig. 2B). This can be attributed to slower hook growth in general, which would result in overall less flagella per cell. The secretion levels of FliK were slightly increased for all FliK_N_ variants in the FlgE_T149N_ background compared to the WT (Fig. 2F). Presumably, the increased FliK secretion is related to the slow-growing hook phenotype, which influences the timing of substrate specificity switching and thus allows additional time for secretion of FliK. Again, FliK_305_ was barely detected in the culture supernatant, which suggested instability of the mutant protein as mentioned above. We next isolated HBB complexes to measure hook-lengths in the short FliK_N_ variants (FliK_385_ to FliK_305_) in the FlgE_T149N_ background. Consistent with our hypothesis that the level of FlgE production, secretion and polymerization contributes to the minimal hook-length, we found shorter hooks of controlled length (i.e. no polyhooks) in the FliK_N_ variants of the slow-hook polymerization background compared to the WT background (Fig. 2H+I and Fig. S6). Interestingly, also the FliK_305_ variant, which displayed uncontrolled hook-length in a WT background, assembled hooks of controlled length in the FlgE_T149N_ background. Slower or inefficient polymerization of the hook likely increases the number of measurements by secreted FliK, resulting in the shorter hook phenotype (3). However, independent from FliK length, hooks shorter than a minimal length of approximately 40 nm were not observed. These results support the hypothesis that FlgE secretion and polymerization speed is an important factor that determines the minimal length of the hook.

### Coupling *flgE* and *fiK* transcription results in shorter hook-length

Based on our observation that the rate of FlgE secretion and polymerization contributes to the hook-length control mechanism, we next devised an alternative strategy to change the ratio of produced and secreted FliK and FlgE. We generated *flgE* and *fliK* transcriptional fusions by introducing the genes either within the *flg* or the *fig* operons in a respective deletion background (Δ*flgE* or Δ*fliK*) downstream of either *fliK* or *flgE*. We constructed these operon fusions using the WT FliK_405_ as well as the short FliK_325_ variant. Motility assays showed a decrease in motility for all four constructs, although the defect was more severe (35-42% of WT) when *flgE* was introduced into the *fli* operon, compared to 65-77% of the WT when *fliK* was introduced into the *flg* operon (Fig. 3A and Fig. S8A).

**Figure 3.**
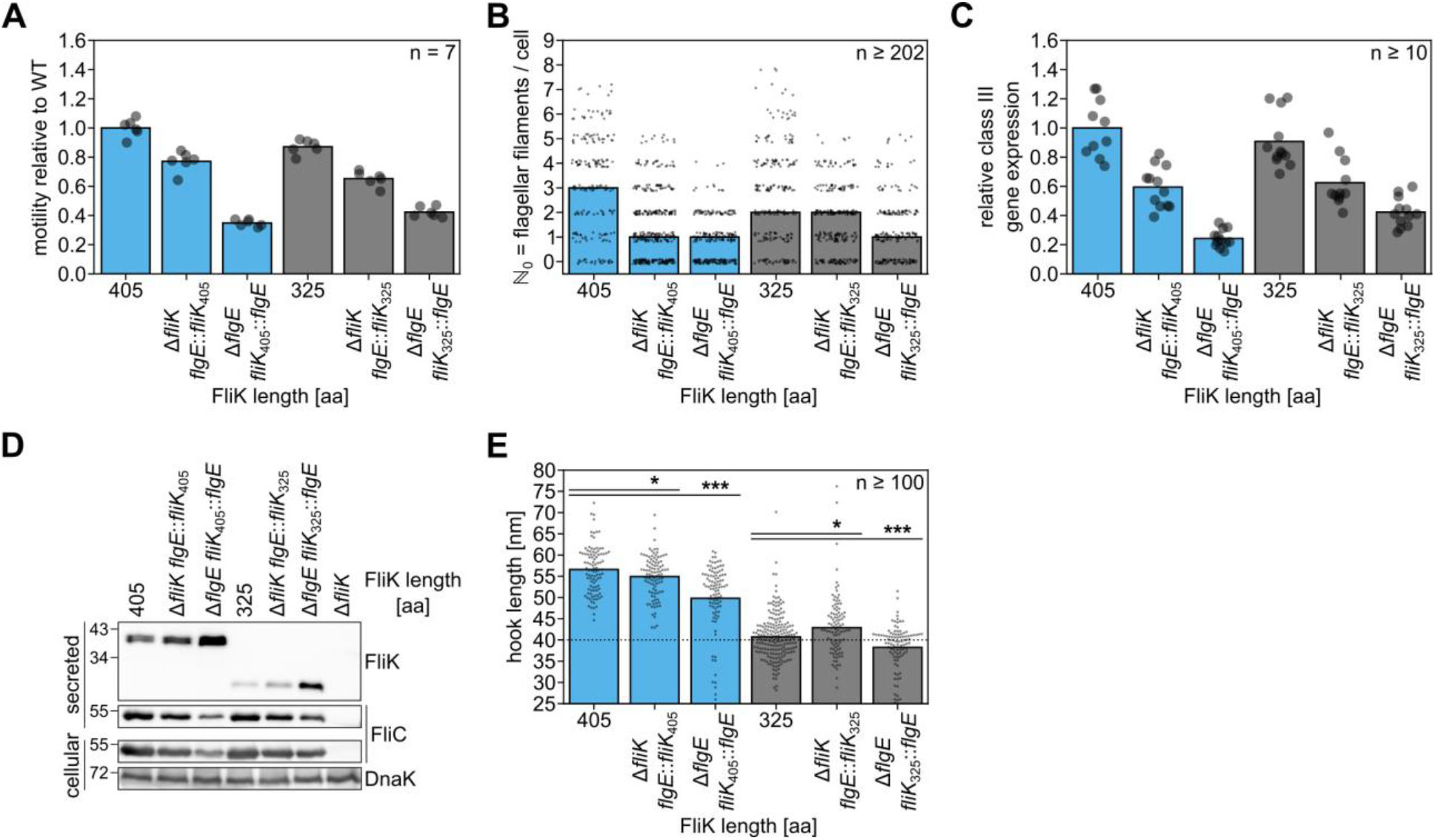
Changing FliK/FlgE expression ratio affects motility, flagellation and hook-length control. (A) Motility phenotype of FliK/FlgE operon fusions (blue, FliK_405 aa_; grey, FliK_325 aa_) analyzed using soft-agar swim plates. Relative motility represents the motility swarm diameters normalized to the WT-FliK_405 aa_. The bar graphs represent the mean of n=7 biologically independent replicates per strain. Replicates are shown as individual data points. (B) Flagellation of FliK/FlgE operon fusions (blue, FliK_405 aa_; grey, FliK_325 aa_) analyzed using anti-flagellin immunostaining. The bar graphs represent the median flagella per cell of n≥202 individual bacteria analyzed per strain. The number of flagella per cell ℕ_0_ are shown as individual data points for each strain. Flagellation data for FliK_405_ and FliK_325_ correspond to those shown in Fig. 2B. (C) Flagellar class 3 gene expression as a reporter of the substrate specificity switch capability in FliK/FlgE operon fusions. Flagellar biosynthesis was synchronized using an anhydrotetracycline-inducible flagellar master regulator (P*_tet_-flhDC*) and analyzed using a P*_motA_-luxCDABE* reporter. The relative class 3 gene expression was determined by measuring luminescence 105 min after induction of FlhDC expression and is reported normalized to the WT strain (FliK_405_). The bar graphs represent the mean of n≥10 biologically independent replicates. Replicates are shown as individual data points. The class 3 gene expression data for FliK_405_ and FliK_325_ correspond to those shown in Fig. 1C. (D) Cytoplasmic and secreted protein levels of FliK and FliC of strains harboring FliK/FlgE operon fusions. DnaK served as loading and cell lysis control. The size of a protein ladder in kDa is indicated on the left. (E) Mean hook-length of FliK/FlgE operon fusions (blue, FliK_405 aa_; grey, FliK_325 aa_). Hook-lengths of n≥100 isolated hook-basal body complexes were determined. Statistical significance was calculated using unpaired Student’s t-test (* = p≤0.05, *** = p≤0.001) The hook-length data for FliK_405_ and FliK_325_ correspond to those shown in Fig. 2G and hook-length distributions for the individual strains are shown in Supporting Information Fig. S8. WT, wildtype; aa, amino acids.

This motility defect correlated with a reduced number of flagella per cell analyzed by immunostaining of the flagellar filaments. We note that we observed a large number of cells without any filaments, suggesting that the timing of the substrate specificity switch was impaired (Fig. 3B and Fig. S8B). In agreement, the estimated time of the substrate specificity switch as determined using the class 3 gene expression reporter was delayed in the operon fusion variants (Fig. 3C). We next determined the cellular and secreted protein levels of FliK and the flagellin FliC in the operon fusion variants (Fig. 3D). When *flgE* was fused to *fliK_405_*, the levels of produced and secreted FliC was reduced, supporting the hypothesis that the hook-basal bodies switched to late substrate secretion. FliK secretion was similar for both fliK_405_ and FliK_325_ when *fliK* was fused to the *flg* operon compared to their respective WT. In contrast, FliK secretion levels were increased when *flgE* was fused to the *fli* operon for both FliK variants. This indicates that during hook assembly of the *fliK-flgE* operon fusions, FlgE was comparatively underproduced. As mentioned above, underproduction of FlgE would result in more frequent FliK measurements of hook-length and, presumably, decrease hook-length. To verify these observations, we purified HBB complexes and determined hook-length. Expressing either fliK_405_ or FliK_325_ from the *flg* operon produced hooks with a similar hook-length distribution to their respective WT background (Fig. 3E and Fig. S8). However, expressing *flgE* from the *fli* operon resulted for both fliK_405_ and FliK_325_ in a shorter hook-length of approximately 50 nm ± 9 nm and 38 nm ± 5 nm, respectively, compared to 56 nm ± 6 nm and 45 ± 8 nm for the WT background, respectively.

## Discussion

The correct self-assembly of the flagellum follows a strict hierarchical process. Presumably to conserve energy and biosynthetic resources, flagellar gene expression is coupled to the assembly state of the growing flagellum and regulated by several distinct mechanisms (1). One of them connects hook-length control with the subsequent substrate specificity switch and expression of late flagellar genes, including flagellin, from class 3 promoters. While the protein components involved in hook-length determination and the substrate specificity switch are known, the underlying mechanism is not completely understood. In particular, the current model of hook-length control predicts that a molecular measuring tape protein, FliK, is intermittently secreted during hook growth and induces the switch in substrate specificity with a certain probability if secretion of FliK has slowed down (3). When the hook is short, FliK is initially secreted fast and thereby unable to induce the substrate specificity switch. As soon as the hook has grown to a certain minimal length (approximately 55 nm in *Salmonella*), the complete FliK molecule remains in the secretion channel for a certain amount of time, which is sufficient for the C-terminal domain of FliK to induce the substrate specificity switch. Because a certain probability exists for a productive interaction of the C-terminal domain of FliK with the export apparatus to induce the substrate specificity switch, this mechanism is presumed to be dependent on the rate of FliK protein secretion. This is supported by observations that overproduction of FliK (or underproduction of the hook subunit FlgE) results in shorter hook-lengths (25–27), whereas underproduction of FliK (or overproduction of FlgE) results in longer hook-lengths (25, 27). The idea that FliK functions as a molecular measuring tape is mainly based on the observation that insertions of α-helical parts of the homologous protein of the *Yersinia* injectisome system into FliK resulted in a linear increase in hook-length by 0.2 nm per inserted amino acid (5, 15). In support, some deletions of the α-helical N-terminal domain of FliK resulted in correspondingly shorter hook-lengths. However, some FliK deletion mutants resulted in hooks of uncontrolled length, which is not consistent with the molecular ruler model (5, 15).

Accordingly, we performed a systematic, stepwise deletion screen of the N-terminal domain of FliK in order to identify and characterize FliK deletion variants that are able to control hook-length. FliK is a 405 aa protein consisting primarily of two domains connected by a flexible linker. The N-terminal domain ranges from aa1-180 with a peptide secretion signal within the first 40 aa, while the C-terminal domain comprises aa206-405 (14). Previously, most functional, shorter FliK variants were constructed with deletions partially within the C-terminal domain and spanning from the N-terminal domain into the linker region (aa181-205) (5, 15, 30, 31). The shortest, functional FliK variant obtained was 363 aa in length (Δaa161-202) and resulted in a hook-length of 44 ± 8 nm (15). By focusing our deletion mutagenesis only on the N-terminal domain of FliK, we were able to construct various deletion variants that displayed only a mild motility defect compared to the WT (Fig. 1A+B and Fig. S1). The extensive deletion analysis further demonstrated the importance of the region aa121-160. Unexpectedly, a deletion of aa121-140 resulted in less motility and slower substrate specificity switching, while longer deletions including this region exhibited partially no or mild defects. We speculate that additional structural or folding properties of the truncated FliK molecule, which depend on the length of the deletion, influence the function of FliK_N_ in hook-length control. Purification of HBB complexes of selected mutants revealed hooks of controlled length (i.e. no polyhooks) up to a FliK length of 325 aa, which is 38 aa shorter than the shortest variant described before (Fig. 2G+I and Fig. S4). However, we surprisingly found that, independent of the length of FliK, the length of the hook did not decrease below a minimal length of approximately 45 nm. Slowing-down hook polymerization by introducing the FlgE_T149N_ mutation into the short FliK variants resulted in the formation of shorter hooks, and thus supports the above-mentioned model that FlgE polymerization speed into the growing hook structure is an important factor for determining the minimal length of the hook (Fig. 2H+I and Fig. S6). However, also in the slow-hook polymerization background FlgE_T149N_, hook-length did not decrease below a minimal length of about 40 nm irrespective of FliK length. The model that the rate of hook polymerization determines the minimal length of the hook is further supported by our analysis of *flgE/fliK* operon fusions. We generated transcriptional fusions of *flgE* and *fliK* in the *fli* or *flg* operons, respectively, with the goal to manipulate the ratio of produced and secreted FlgE vs. FliK. Transcription of *flgE* is approximately 2-fold higher than *fliK* transcription in the WT (32). It is therefore reasonable that *flgE* transcription might be reduced when fused to *fliK* in the *fli* operon. Accordingly, a decreased FlgE secretion and polymerization compared to the WT resulted in shorter hooks in the fliK-flgE operon fusion due to comparatively more frequent secretion of FliK (Fig. 3D+E) (3). Unexpectedly, re-organizing the gene order in the *fli* and *flg* operons led to a severe defect in motility and a reduction in the number of flagella per cell, although hook-length was controlled (Fig. 3A+B+E). This result can be attributed to the slower substrate specificity switching compared to the WT and highlights the significance of proper coupling of gene expression during the hierarchical assembly of the flagellum (Fig. 3C).

In summary, the present work describes the shortest FliK variants reported so far that are able to control hook-length. However, the short FliK variants were not able to control hook-length below a certain threshold, which is not consistent with a molecular ruler mechanism, where FliK physically measures the correct hook-length. We found that the minimal length of the hook rather depends on the level of produced and secreted hook subunit FlgE and thus on the rate of hook growth. This is supported by a previous growth rate analysis of polyhooks, which showed that the hook elongation rate is initially fast (40 nm/min) and slows to 8 nm/min for hooks greater than 55 nm (33).

Accordingly, our results support a model in which FliK functions as a hook growth terminator, which prevents the hook from growing beyond the length required for its universal joint function (34). In this model, both hook and FliK subunits are secreted during hook assembly. Initially, the rate of hook subunit secretion is fast and likely few FliK molecules will be exported. In addition, it has been shown that FliK_N_ folds into a ball-like shape (35) and thus, when the hook is still short, FliK_N_ will start folding as soon as it exits the hook tip. The folding of FliK_N_ will rapidly pull the rest of FliK outside of the cell, and thus prevent an interaction of the C-terminal domain of FliK with the FlhB component of the T3SS for short hooks (3, 36). The longer the hook gets, the slower its growth will be and if another FliK molecule is secreted in a hook with a length of approximately 55 nm or longer, FliK_N_ will remain in the channel for some time as α-helical polypeptide before exiting the hook tip. This allows sufficient time for an interaction of the C-terminal domain of FliK with FlhB in order to switch the secretion substrate specificity and therefore end further hook growth. The hook growth terminator FliK thus controls the maximal, but not the minimal length of the flagellar hook.

## Materials and Methods

### Bacterial strains, plasmids and media

All bacterial strains used in this study are listed in Supplemental Table S1. Cells were grown in lysogeny broth (LB). Motility agar (0.3%) was prepared based on TB broth (1% tryptone and 0.5% NaCl). The generalized transducing phage of *Salmonella enterica* serovar Typhimurium P22 *HT105*/*1 int-201* was used in all transductional crosses (37). Deletions were constructed using λ-RED homologous recombination (38, 39). Medium was supplemented with 12.5 μg/ml chloramphenicol (Cm) or 100 ng/ml anhydrotetracycline (AnTc) if necessary.

### Motility assay

Motility of bacteria was determined by inoculating 1 μl overnight culture into semisolid agar plates (0.3%). Plates were incubated for 4.5 h at 37 °C or 8 h at 30 °C. Swimming halos were analyzed using ImageJ and the obtained values were normalized to the respective WT (fliK_405_ or fliK_405_ FlgE_T149N_).

### Secretion assay

An overnight culture of FliC-phase locked bacteria (Δ*hin*-5717) was diluted 1:100 in LB broth and incubated at 37 °C or 30 °C until the bacteria reached late exponential phase. 1.9 ml culture was harvested and supernatant and pellet fractions were separated by centrifugation (3 min, 13,000 × g, 4°C). Proteins were precipitated using a final concentration of 10% TCA. Samples were resuspended in SDS sample buffer normalized to the OD600. 200 OD units were loaded for SDS-PAGE and subsequent western blotting. Proteins of interest were detected using primary polyclonal anti-FliK and anti-FliC (BD) antibodies as well as monoclonal anti-DnaK (abcam) antibodies and secondary antibodies (anti-rabbit, anti-mouse) coupled to HRP.

### Immunostaining of flagellar filaments and fluorescent microscopy

An overnight culture of FliC-phase locked bacteria (Δ*hin*-5717) was diluted 1:100 in LB broth and incubated at 37 °C or 30 °C until the bacteria reached mid-log phase. Bacteria were fixed with PFA at a final concentration of 2% for 10 min at room temperature and applied to a homemade flow-cell (40). Cells were blocked for 10 min with 10% BSA. The primary anti-FliC antibody (BD) was added for 1 h at room temperature. The secondary anti-rabbit Alexa Fluor 488 antibody (Thermo Fisher Scientific) was incubated for 30 min at room temperature. The slide was washed twice with 1×PBS and bacteria were mounted with Fluoroshield™ with DAPI (Sigma). Bacteria were observed with a Zeiss Axio Observer Z1 inverted microscope and images were acquired at 100× magnification with an Axiocam 506 mono CCD-camera using the Zen 2.6 pro software. Individual flagella were counted manually using ImageJ.

### Purification of hook-basal body complexes

Hook-basal body complexes were purified as described before (5, 41). Purified samples were stored at 4 °C with a final concentration of 1% formaldehyde and shipped to Manfred Rohde at the Helmholtz Centre for Infection Research, Braunschweig, Germany, for imaging and hook-length measurements.

### Negative staining for transmission electron microscopy

Thin carbon support films were prepared by sublimation of a carbon thread onto a freshly cleaved mica surface. The different hook-length preparations were absorbed onto the carbon film, washed in TE buffer (20 mM Tris, 2 mM EDTA, pH 7.0) and distilled water. Adsorbed hooks were negatively stained with 2% aqueous uranyl acetate, pH 5.0. After air-drying, samples were examined in an EM 910 transmission electron microscope (Zeiss) at an acceleration voltage of 80 kV. Images were taken at calibrated magnifications using a line replica. Images were recorded digitally with a Slow-Scan CCD-Camera (ProScan, 1024×1024) applying the ITEM-Software (Olympus Soft Imaging Solutions).

### Luciferase assay

For analyzing the substrate specificity switching capability and timing the plasmid pRG19 (P*_motA_*-*luxCDABE*) was used as a class 3 reporter (21, 22). Single colonies were inoculated in 200 μl LB supplemented with 12.5 μg/ml Cm in a 96-well plate and incubated for 8 h at 37 °C with agitation. Bacteria were diluted 1:100 in LB with Cm and AnTc at a final concentration of 100 ng/ml in a white 96-well plate to induce transcription of the flagellar master regulator (P*_tetA_*-*flhDC*) (23). Luminescence and OD_600_ was measured over 6 h in 15 min intervals using a Synergy H1 microplate reader (BioTek). Relative light units (RLU) were calculated as:

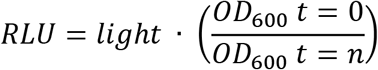

Values were normalized to the WT.

## Acknowledgments

We thank Heidi Landmesser for technical assistance, Marina Douanla for help in strain construction, Ina Schleicher for assistance in TEM experiments, Tohru Minamino for the gift of anti-FliK antibodies, Kelly Hughes for generous donation of strains and Shin-Ichi Aizawa, as well as members of the Erhardt lab for discussions and helpful comments on the manuscript. This work was supported by start-up funds of the Humboldt-Universität zu Berlin (to M.E.). The funders had no role in study design, data collection and analysis, decision to publish, or preparation of the manuscript.

## Author contributions

A.G. and M.E. conceived the study and designed research; A.G. and M.R. performed experiments; A.G., M.R. and M.E. analyzed data; A.G., M.E. and M.R. wrote the paper; M.R. and M.E. contributed funding and resources.

