## Supporting Information for "Controlling minimal and maximal hook-length of the bacterial flagellum"

**Supplemental Table S1:** List of *Salmonella enterica* serovar Typhimurium LT2 strains used in this study.

| Strain | Genotype | Reference |
|---|---|---|
| TH437 | LT2 wildtype – $FliK_{405}$ | John Roth |
| EM7926 | Δ*fliK*23066 (Δaa41-60) – $FliK_{385}$ | This study |
| EM7927 | Δ*fliK*23067 (Δaa61-80) | This study |
| EM7928 | Δ*fliK*23068 (Δaa81-100) | This study |
| EM7929 | Δ*fliK*23069 (Δaa101-120) | This study |
| EM7930 | Δ*fliK*23070 (Δaa121-140) | This study |
| EM7931 | Δ*fliK*23071 (Δaa141-160) | This study |
| EM7932 | Δ*fliK*23072 (Δaa161-180) | This study |
| EM8047 | Δ*fliK*23086 (Δaa61-100) | This study |
| EM8048 | Δ*fliK*23087 (Δaa81-120) | This study |
| EM8049 | Δ*fliK*23088 (Δaa101-140) | This study |
| EM8050 | Δ*fliK*23089 (Δaa121-160) | This study |
| EM8051 | Δ*fliK*23090 (Δaa141-180) | This study |
| EM8263 | Δ*fliK*23129 (Δaa41-80) – $FliK_{365}$ | This study |
| EM8264 | Δ*fliK*23130 (Δaa61-120) | This study |
| EM8265 | Δ*fliK*23131 (Δaa81-140) | This study |
| EM8266 | Δ*fliK*23132 (Δaa101-160) | This study |
| EM8267 | Δ*fliK*23133 (Δaa41-180) – $FliK_{265}$ | This study |
| EM8277 | Δ*fliK*23134 (Δaa41-100) – $FliK_{345}$ | This study |
| EM8279 | Δ*fliK*23135 (Δaa121-180) | This study |
| EM8280 | Δ*fliK*23136 (Δaa41-120) – $FliK_{325}$ | This study |
| EM8281 | Δ*fliK*23137 (Δaa61-140) | This study |
| EM8282 | Δ*fliK*23138 (Δaa81-160) | This study |
| EM8283 | Δ*fliK*23139 (Δaa101-180) | This study |
| EM8284 | Δ*fliK*23140 (Δaa41-140) – $FliK_{305}$ | This study |
| EM8285 | Δ*fliK*23141 (Δaa81-180) | This study |
| EM8321 | Δ*fliK*23143 (Δaa61-160) | This study |
| EM8545 | Δ*fliK*23066 (Δaa41-60) Δ*hin*-5717::FCF (*fliC*$^{ON}$) – $FliK_{385}$ | This study |
| EM8546 | Δ*fliK*23067 (Δaa61-80) Δ*hin*-5717::FCF (*fliC*$^{ON}$) | This study |
| EM8547 | Δ*fliK*23068 (Δaa81-100) Δ*hin*-5717::FCF (*fliC*$^{ON}$) | This study |
| EM8548 | Δ*fliK*23069 (Δaa101-120) Δ*hin*-5717::FCF (*fliC*$^{ON}$) | This study |
| EM8549 | Δ*fliK*23070 (Δaa121-140) Δ*hin*-5717::FCF (*fliC*$^{ON}$) | This study |
| EM8550 | Δ*fliK*23071 (Δaa141-160) Δ*hin*-5717::FCF (*fliC*$^{ON}$) | This study |
| EM8551 | Δ*fliK*23072 (Δaa161-180) Δ*hin*-5717::FCF (*fliC*$^{ON}$) | This study |
| EM8552 | Δ*fliK*23129 (Δaa41-80) Δ*hin*-5717::FCF (*fliC*$^{ON}$) – $FliK_{365}$ | This study |

| | | |
|---|---|---|
| EM8553 | Δ*fliK*23086 (Δaa61-100) Δ*hin*-5717::FCF (*fliC*$^{ON}$) | This study |
| EM8554 | Δ*fliK*23087 (Δaa81-120) Δ*hin*-5717::FCF (*fliC*$^{ON}$) | This study |
| EM8555 | Δ*fliK*23088 (Δaa101-140) Δ*hin*-5717::FCF (*fliC*$^{ON}$) | This study |
| EM8556 | Δ*fliK*23089 (Δaa121-160) Δ*hin*-5717::FCF (*fliC*$^{ON}$) | This study |
| EM8557 | Δ*fliK*23090 (Δaa141-180) Δ*hin*-5717::FCF (*fliC*$^{ON}$) | This study |
| EM8558 | Δ*fliK*23134 (Δaa41-100) Δ*hin*-5717::FCF (*fliC*$^{ON}$) – $FliK_{345}$ | This study |
| EM8559 | Δ*fliK*23130 (Δaa61-120) Δ*hin*-5717::FCF (*fliC*$^{ON}$) | This study |
| EM8560 | Δ*fliK*23131 (Δaa81-140) Δ*hin*-5717::FCF (*fliC*$^{ON}$) | This study |
| EM8561 | Δ*fliK*23132 (Δaa101-160) Δ*hin*-5717::FCF (*fliC*$^{ON}$) | This study |
| EM8562 | Δ*fliK*23135 (Δaa121-180) Δ*hin*-5717::FCF (*fliC*$^{ON}$) | This study |
| EM8563 | Δ*fliK*23136 (Δaa41-120) Δ*hin*-5717::FCF (*fliC*$^{ON}$) – $FliK_{325}$ | This study |
| EM8564 | Δ*fliK*23137 (Δaa61-140) Δ*hin*-5717::FCF (*fliC*$^{ON}$) | This study |
| EM8565 | Δ*fliK*23138 (Δaa81-160) Δ*hin*-5717::FCF (*fliC*$^{ON}$) | This study |
| EM8566 | Δ*fliK*23139 (Δaa101-180) Δ*hin*-5717::FCF (*fliC*$^{ON}$) | This study |
| EM8567 | Δ*fliK*23140 (Δaa41-140) Δ*hin*-5717::FCF (*fliC*$^{ON}$) – $FliK_{305}$ | This study |
| EM8568 | Δ*fliK*23143 (Δaa61-160) Δ*hin*-5717::FCF (*fliC*$^{ON}$) | This study |
| EM8569 | Δ*fliK*23141 (Δaa81-180) Δ*hin*-5717::FCF (*fliC*$^{ON}$) | This study |
| EM8570 | Δ*fliK*23133 (Δaa41-180) Δ*hin*-5717::FCF (*fliC*$^{ON}$) – $FliK_{265}$ | This study |
| EM8586 | Δ*fliK*23066 (Δaa41-60) $P_{flhDC5451}$::Tn*10d*Tc[del-25] / pRG19::FCF ($P_{motA}$-*luxCDABE*, $Cm^R$ $Tet^R$) – $FliK_{385}$ | This study |
| EM8587 | Δ*fliK*23067 (Δaa61-80) $P_{flhDC5451}$::Tn*10d*Tc[del-25] / pRG19::FCF ($P_{motA}$-*luxCDABE*, $Cm^R$ $Tet^R$) | This study |
| EM8588 | Δ*fliK*23068 (Δaa81-100) $P_{flhDC5451}$::Tn*10d*Tc[del-25] / pRG19::FCF ($P_{motA}$-*luxCDABE*, $Cm^R$ $Tet^R$) | This study |
| EM8589 | Δ*fliK*23069 (Δaa101-120) $P_{flhDC5451}$::Tn*10d*Tc[del-25] / pRG19::FCF ($P_{motA}$-*luxCDABE*, $Cm^R$ $Tet^R$) | This study |
| EM8590 | Δ*fliK*23070 (Δaa121-140) $P_{flhDC5451}$::Tn*10d*Tc[del-25] / pRG19::FCF ($P_{motA}$-*luxCDABE*, $Cm^R$ $Tet^R$) | This study |
| EM8591 | Δ*fliK*23071 (Δaa141-160) $P_{flhDC5451}$::Tn*10d*Tc[del-25] / pRG19::FCF ($P_{motA}$-*luxCDABE*, $Cm^R$ $Tet^R$) | This study |
| EM8592 | Δ*fliK*23072 (Δaa161-180) $P_{flhDC5451}$::Tn*10d*Tc[del-25] / pRG19::FCF ($P_{motA}$-*luxCDABE*, $Cm^R$ $Tet^R$) | This study |
| EM8593 | Δ*fliK*23129 (Δaa41-80) $P_{flhDC5451}$::Tn*10d*Tc[del-25] / pRG19::FCF ($P_{motA}$-*luxCDABE*, $Cm^R$ $Tet^R$) – $FliK_{365}$ | This study |
| EM8594 | Δ*fliK*23086 (Δaa61-100) $P_{flhDC5451}$::Tn*10d*Tc[del-25] / pRG19::FCF ($P_{motA}$-*luxCDABE*, $Cm^R$ $Tet^R$) | This study |
| EM8595 | Δ*fliK*23087 (Δaa81-120) $P_{flhDC5451}$::Tn*10d*Tc[del-25] / pRG19::FCF ($P_{motA}$-*luxCDABE*, $Cm^R$ $Tet^R$) | This study |
| EM8596 | Δ*fliK*23088 (Δaa101-140) $P_{flhDC5451}$::Tn*10d*Tc[del-25] / pRG19::FCF ($P_{motA}$-*luxCDABE*, $Cm^R$ $Tet^R$) | This study |
| EM8597 | Δ*fliK*23089 (Δaa121-160) $P_{flhDC5451}$::Tn*10d*Tc[del-25] / pRG19::FCF ($P_{motA}$-*luxCDABE*, $Cm^R$ $Tet^R$) | This study |
| EM8598 | Δ*fliK*23090 (Δaa141-180) $P_{flhDC5451}$::Tn*10d*Tc[del-25] / pRG19::FCF ($P_{motA}$-*luxCDABE*, $Cm^R$ $Tet^R$) | This study |

| | | |
|---|---|---|
| EM8599 | Δ*fliK*6140 (Δaa6-end) $P_{flhDC5451}$::Tn*10d*Tc[del-25] / pRG19::FCF ($P_{motA}$-*luxCDABE*, $Cm^R$ $Tet^R$) | This study |
| EM8644 | Δ*fliK*23134 (Δaa41-100) $P_{flhDC5451}$::Tn*10d*Tc[del-25] / pRG19::FCF ($P_{motA}$-*luxCDABE*, $Cm^R$ $Tet^R$) – $FliK_{345}$ | This study |
| EM8645 | Δ*fliK*23130 (Δaa61-120) $P_{flhDC5451}$::Tn*10d*Tc[del-25] / pRG19::FCF ($P_{motA}$-*luxCDABE*, $Cm^R$ $Tet^R$) | This study |
| EM8646 | Δ*fliK*23131 (Δaa81-140) $P_{flhDC5451}$::Tn*10d*Tc[del-25] / pRG19::FCF ($P_{motA}$-*luxCDABE*, $Cm^R$ $Tet^R$) | This study |
| EM8647 | Δ*fliK*23132 (Δaa101-160) $P_{flhDC5451}$::Tn*10d*Tc[del-25] / pRG19::FCF ($P_{motA}$-*luxCDABE*, $Cm^R$ $Tet^R$) | This study |
| EM8648 | Δ*fliK*23135 (Δaa121-180) $P_{flhDC5451}$::Tn*10d*Tc[del-25] / pRG19::FCF ($P_{motA}$-*luxCDABE*, $Cm^R$ $Tet^R$) | This study |
| EM8649 | Δ*fliK*230136 (Δaa41-120) $P_{flhDC5451}$::Tn*10d*Tc[del-25] / pRG19::FCF ($P_{motA}$-*luxCDABE*, $Cm^R$ $Tet^R$) – $FliK_{325}$ | This study |
| EM8650 | Δ*fliK*23137 (Δaa61-140) $P_{flhDC5451}$::Tn*10d*Tc[del-25] / pRG19::FCF ($P_{motA}$-*luxCDABE*, $Cm^R$ $Tet^R$) | This study |
| EM8651 | Δ*fliK*23138 (Δaa81-160) $P_{flhDC5451}$::Tn*10d*Tc[del-25] / pRG19::FCF ($P_{motA}$-*luxCDABE*, $Cm^R$ $Tet^R$) | This study |
| EM8652 | Δ*fliK*23139 (Δaa101-180) $P_{flhDC5451}$::Tn*10d*Tc[del-25] / pRG19::FCF ($P_{motA}$-*luxCDABE*, $Cm^R$ $Tet^R$) | This study |
| EM8653 | Δ*fliK*23140 (Δaa41-140) $P_{flhDC5451}$::Tn*10d*Tc[del-25] / pRG19::FCF ($P_{motA}$-*luxCDABE*, $Cm^R$ $Tet^R$) – $FliK_{305}$ | This study |
| EM8654 | Δ*fliK*23143 (Δaa61-160) $P_{flhDC5451}$::Tn*10d*Tc[del-25] / pRG19::FCF ($P_{motA}$-*luxCDABE*, $Cm^R$ $Tet^R$) | This study |
| EM8655 | Δ*fliK*23141 (Δaa81-180) $P_{flhDC5451}$::Tn*10d*Tc[del-25] / pRG19::FCF ($P_{motA}$-*luxCDABE*, $Cm^R$ $Tet^R$) | This study |
| EM8656 | Δ*fliK*23133 (Δaa41-180) $P_{flhDC5451}$::Tn*10d*Tc[del-25] / pRG19::FCF ($P_{motA}$-*luxCDABE*, $Cm^R$ $Tet^R$) – $FliK_{265}$ | This study |
| EM8745 | Δ*fliK*23159 (Δaa31-40) | This study |
| EM8746 | Δ*fliK*23160 (Δaa21-40) | This study |
| EM8747 | Δ*fliK*23161 (Δaa41-160) – $FliK_{285}$ | This study |
| EM8809 | Δ*fliK*23161 (Δaa41-160) Δ*hin*-5717::FCF – $FliK_{285}$ | This study |
| EM8833 | Δ*fliK*23161 (Δaa41-160) $P_{flhDC5451}$::Tn*10d*Tc[del-25] / pRG19::FCF ($P_{motA}$-*luxCDABE*, $Cm^R$ $Tet^R$) – $FliK_{285}$ | This study |
| TH14053 | *flgE*2219 (T149N) – $FliK_{405}$ | Lab collection |
| EM9122 | Δ*fliK*23066 (Δaa41-60) *flgE*2219 (T149N) – $FliK_{385}$ | This study |
| EM9123 | Δ*fliK*23129 (Δaa41-80) *flgE*2219 (T149N) – $FliK_{365}$ | This study |
| EM9124 | Δ*fliK*23134 (Δaa41-100) *flgE*2219 (T149N) – $FliK_{345}$ | This study |
| EM9125 | Δ*fliK*23136 (Δaa41-120) *flgE*2219 (T149N) – $FliK_{325}$ | This study |
| EM9176 | *flgE*2219 (T149N) Δ*hin*-5717::FCF ($fliC^{ON}$) | This study |
| EM9171 | Δ*fliK*23066 (Δaa41-60) *flgE*2219 (T149N) Δ*hin*-5717::FCF – $FliK_{385}$ | This study |
| EM9172 | Δ*fliK*23129 (Δaa41-80) *flgE*2219 (T149N) Δ*hin*-5717::FCF – $FliK_{365}$ | This study |

| | | |
|---|---|---|
| EM9173 | Δ*fliK*23134 (Δaa41-100) *flgE*2219 (T149N) Δ*hin*-5717::FCF – $FliK_{345}$ | This study |
| EM9174 | Δ*fliK*23136 (Δaa41-120) *flgE*2219 (T149N) Δ*hin*-5717::FCF – $FliK_{325}$ | This study |
| EM9227 | Δ*fliK*6140 (Δaa6-end) *flgE*23199::*fliK* (full length; operon fusion) | This study |
| EM9228 | Δ*fliK*6140 (Δaa6-end) *flgE*23200::*fliK* (Δaa41-120; operon fusion) | This study |
| EM9281 | Δ*flgE*7599 *fliK*23204 (full length)::*flgE* (after stop; operon fusion) | This study |
| EM9282 | Δ*flgE*7599 *fliK*23205 (Δaa41-120)::*flgE* (after stop; operon fusion) | This study |
| EM9343 | Δ*fliK*6140 (Δaa6-end) *flgE*23199::*fliK* (full length; operon fusion) Δ*hin*-5717::FCF (*fliC*$^{ON}$) | This study |
| EM9344 | Δ*fliK*6140 (Δaa6-end) *flgE*23200::*fliK* (Δaa41-120; operon fusion) Δ*hin*-5717::FCF (*fliC*$^{ON}$) | This study |
| EM9345 | Δ*flgE*7599 *fliK*23204 (full length)::*flgE* (after stop; operon fusion) Δ*hin*-5717::FCF (*fliC*$^{ON}$) | This study |
| EM9346 | Δ*flgE*7599 *fliK*23205 (Δaa41-120)::*flgE* (after stop; operon fusion) Δ*hin*-5717::FCF (*fliC*$^{ON}$) | This study |
| EM9380 | Δ*fliK*23140 (Δaa41-140) *flgE*2219 (T149N) – $FliK_{305}$ | This study |
| EM9673 | Δ*fliK*23140 (Δaa41-140) *flgE*2219 (T149N) Δ*hin*-5717::FCF (*fliC*$^{ON}$) – $FliK_{305}$ | This study |
| TH5633 | $P_{flhDC5451}$::Tn*10d*Tc[del-25] / pRG19::FCF ($P_{motA}$-*luxCDABE*, $Cm^R$ $Tet^R$) – $FliK_{405}$ | Lab collection |
| TH13452 | Δ*fliK*6140 (Δaa6-end) | Lab collection |
| TH14300 | Δ*fliK*6140 (Δaa6-end) Δ*hin*-5717::FRT | Lab collection |
| TH5861 | Δ*hin*-5717::FCF (*fliC*$^{ON}$) – $FliK_{405}$ | Lab collection |
| EM9831 | Δ*fliK*6140 (Δaa6-end) *flgE*23199::*fliK* (full length; operon fusion) $P_{flhDC5451}$::Tn*10d*Tc[del-25] / pRG19::FCF pRG19::FCF ($P_{motA}$-*luxCDABE*, $Cm^R$ $Tet^R$) | This study |
| EM9832 | Δ*fliK*6140 (Δaa6-end) *flgE*23200::*fliK* (Δaa41-120; operon fusion) $P_{flhDC5451}$::Tn*10d*Tc[del-25] / pRG19::FCF pRG19::FCF ($P_{motA}$-*luxCDABE*, $Cm^R$ $Tet^R$) | This study |
| EM9833 | Δ*flgE*7599 *fliK*23204 (full length)::*flgE* (after stop; operon fusion) $P_{flhDC5451}$::Tn*10d*Tc[del-25] / pRG19::FCF (pRG19::FCF ($P_{motA}$-*luxCDABE*, $Cm^R$ $Tet^R$) | This study |
| EM9834 | Δ*flgE*7599 *fliK*23205 (Δaa41-120)::*flgE* (after stop; operon fusion) $P_{flhDC5451}$::Tn*10d*Tc[del-25] / pRG19::FCF ($P_{motA}$-*luxCDABE*, $Cm^R$ $Tet^R$) | This study |

### Supplemental Figures

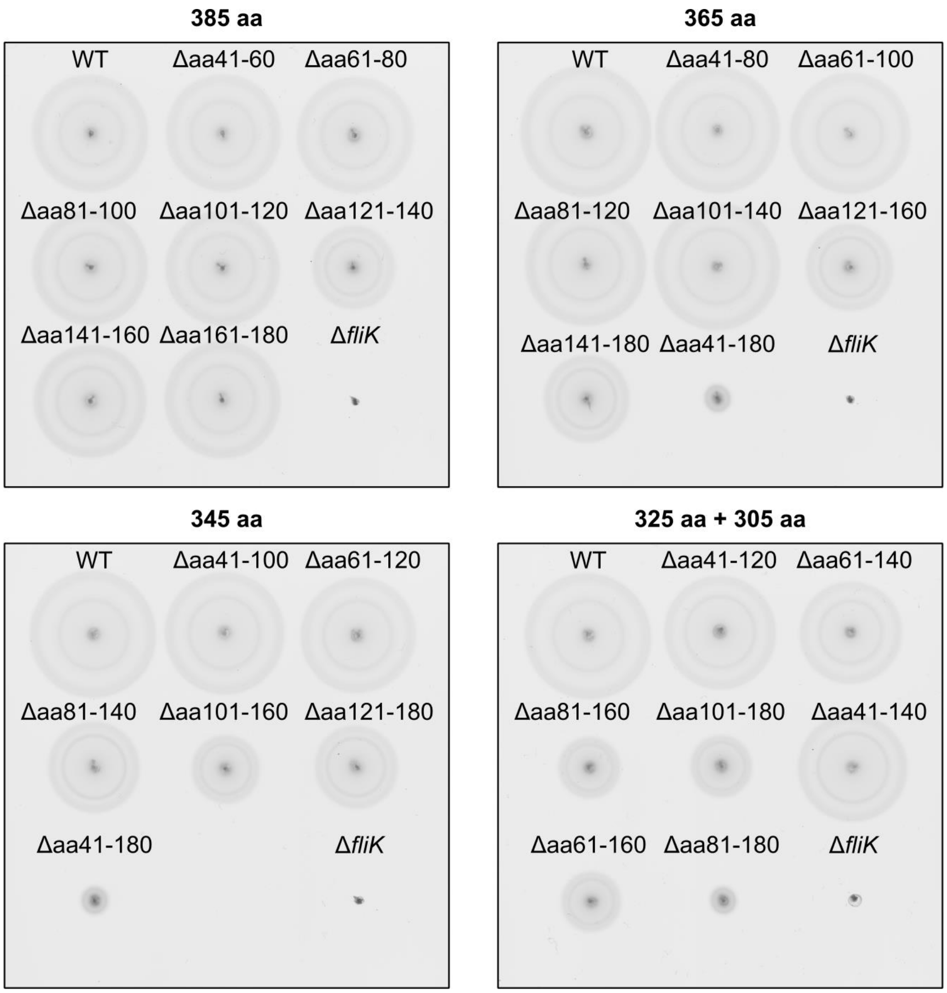


**Supplemental Figure S1. Motility of FliK$_N$ deletion mutants.** Representative soft-agar (0.3%) motility plates of the WT (FliK$_{405\ aa}$) and FliK$_N$ deletion variants incubated for 4.5 h at 37 °C. Deletions within the N-terminal domain of FliK and resulting lengths of the truncated FliK variants are indicated. WT, wildtype; aa, amino acids.

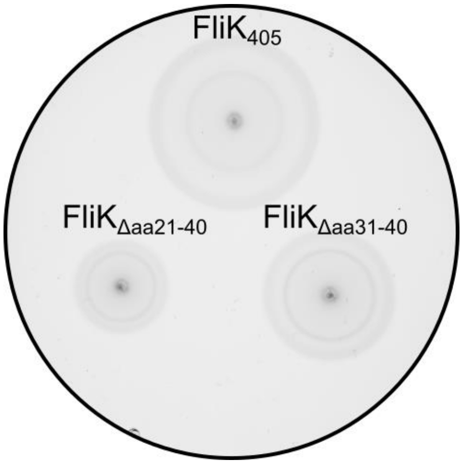


**Supplemental Figure S2. Motility of FliK$_N$ mutants deleted for part of the N-terminal secretion signal.** A representative soft-agar (0.3%) motility plate incubated for 4.5 h at 37 °C is shown. aa, amino acid.

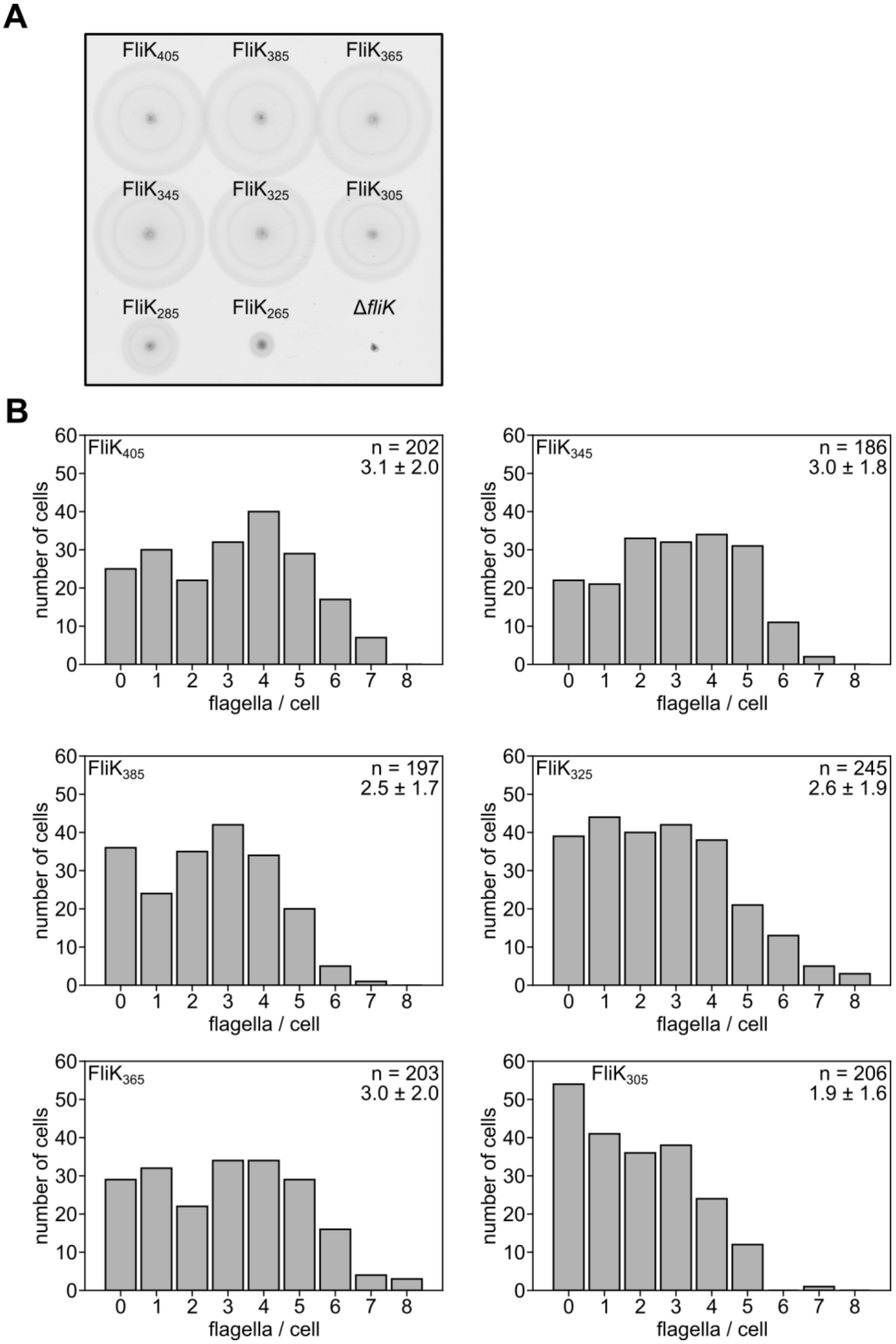


**Supplemental Figure S3. Motility and flagellation phenotype of selected FliK$_{N}$ variants in the WT background.** (A) Representative soft-agar (0.3%) motility plate incubated for 4.5 h at 37 °C. (B) Distribution of flagellar filaments per cell of the WT (FliK$_{405}$) and selected FliK$_{N}$ variants analyzed using anti-flagellin immunostaining. The number of analyzed, individual bacteria (n) and the average number of flagella per cell ± SD is indicated. WT, wildtype; SD, standard deviation.

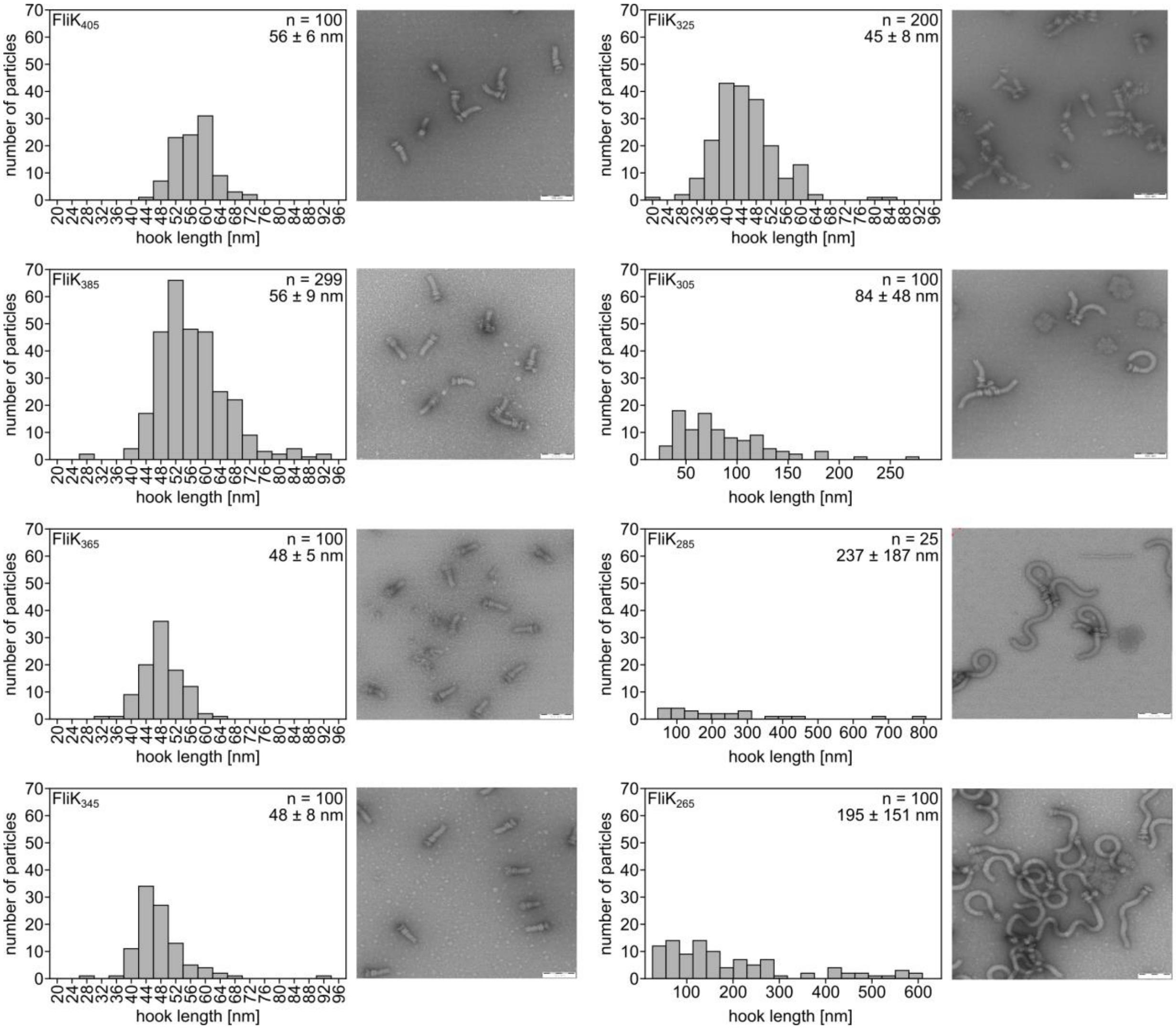


**Supplemental Figure S4. Hook-lengths of selected FliK$_{N}$ variants in the WT background.** Left panels: Hook-length distribution of purified HBBs of the WT (FliK$_{405}$) and selected FliK$_{N}$ variants. The number of measured HBBs (n) and average hook-length ± SD is indicated. Right panels: Representative electron micrographs of purified HBBs. Scale bar = 100 nm. WT, wildtype; HBB, hook-basal body; SD, standard deviation.

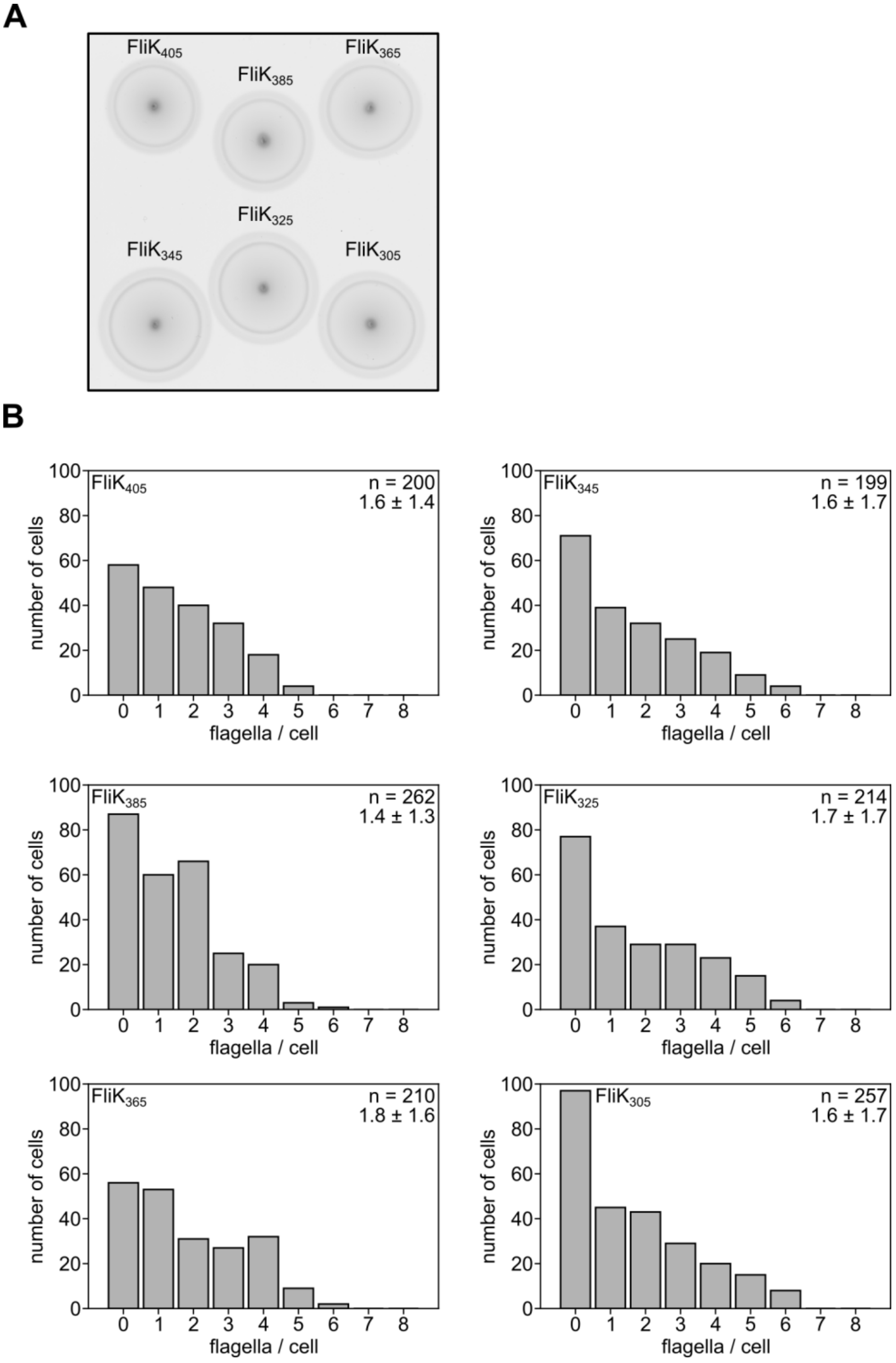


**Supplemental Figure S5. Motility and flagellation phenotype of selected FliK$_{N}$ mutants in FlgE$_{T149N}$ background.** (A) Representative soft-agar (0.3%) motility plate incubated for 8 h at 30 °C. (B) Distribution of flagellar filaments per cell of the WT (FliK$_{405}$) and FliK$_{N}$ variants analyzed using anti-flagellin immunostaining. The number of analyzed, individual bacteria (n) and the average number of flagella per cell ± SD is indicated. WT, wildtype; SD, standard deviation.

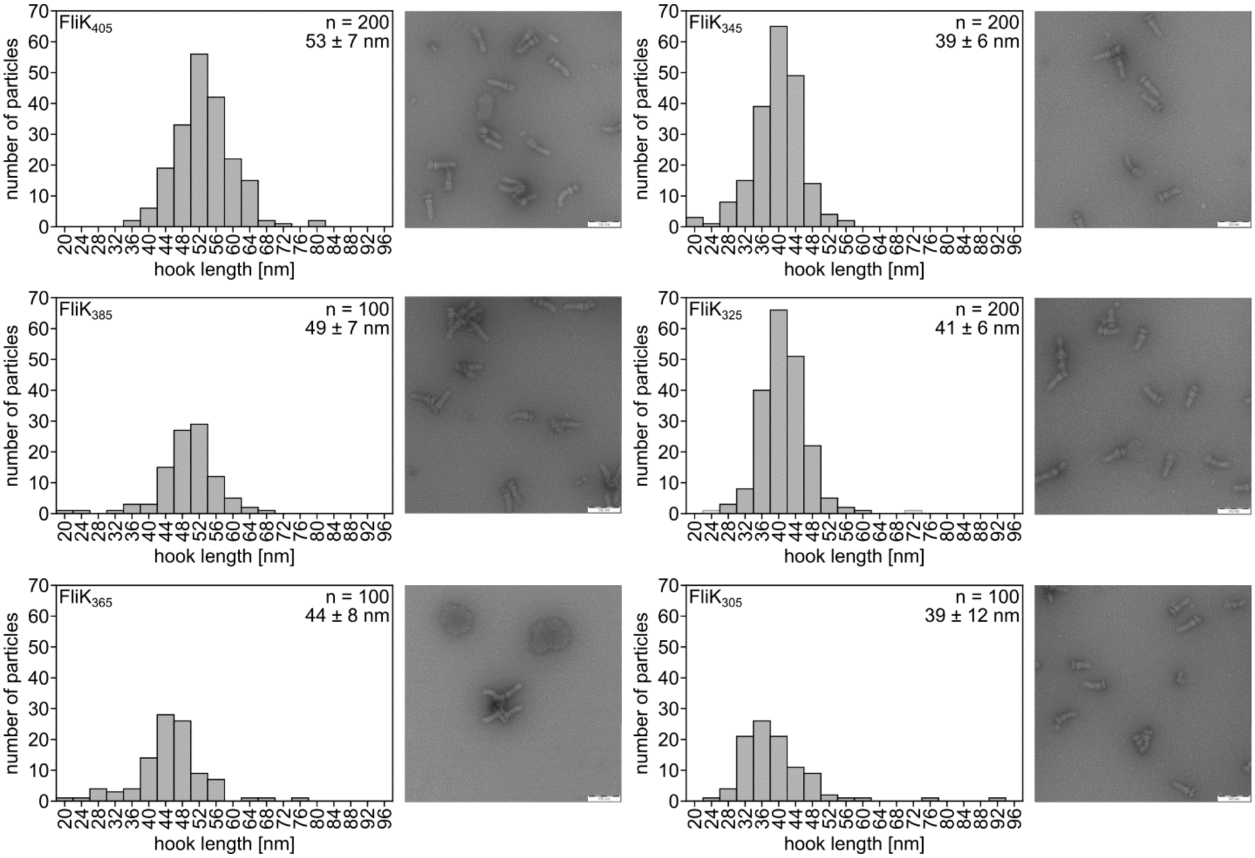


**Supplemental Figure S6. Hook-lengths of selected FliK$_{N}$ variants in the slow-hook polymerization mutant FlgE$_{T149N}$.** Left panels: Hook-length distribution of purified HBBs of the WT (FliK$_{405}$) and selected FliK$_{N}$ variants in the slow-hook polymerization background (FlgE$_{T149N}$). The number of measured HBBs (n) and average hook-length ± SD is indicated. Right panels: Representative electron micrographs of purified HBBs. Scale bar = 100 nm. WT, wildtype; HBB, hook-basal body; SD, standard deviation.

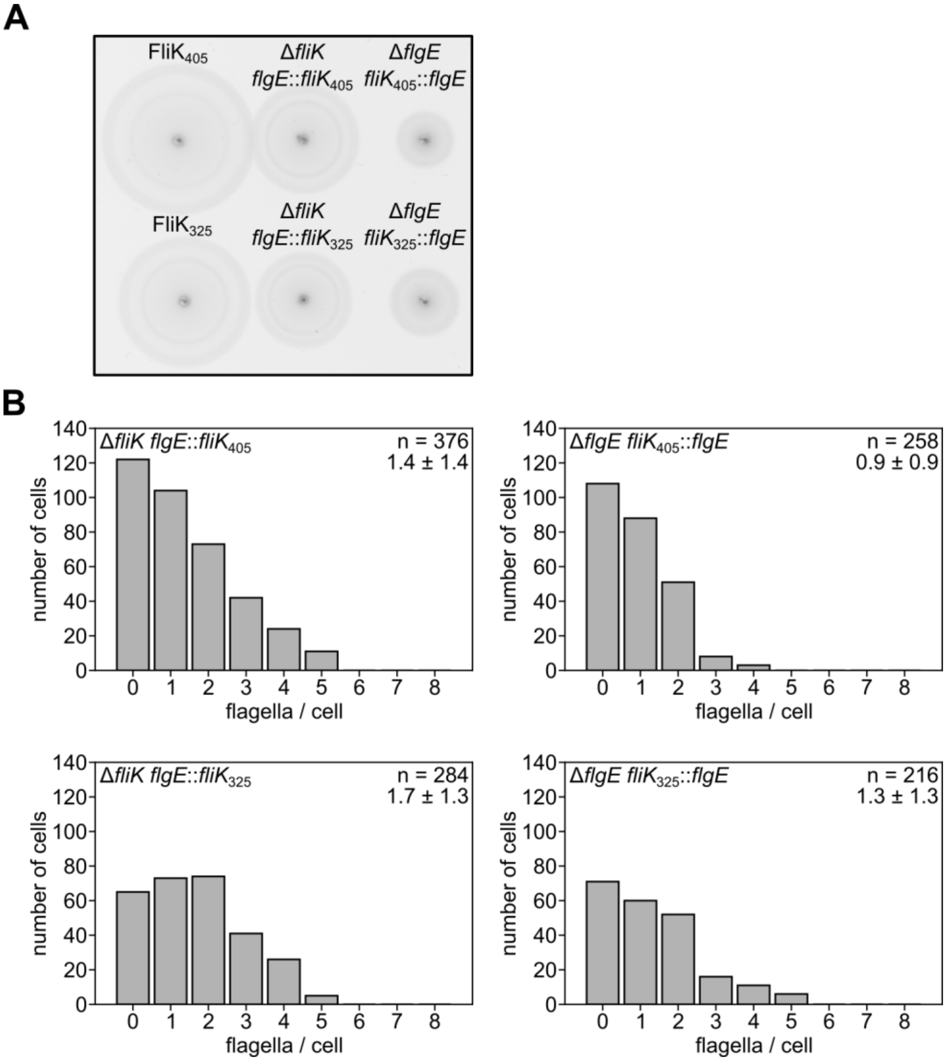


**Supplemental Figure S7. Motility and flagellation pattern of FliK/FlgE operon fusions.** (A) Representative soft-agar (0.3%) motility plate incubated for 4.5 h at 37 °C. (B) Distribution of flagellar filaments per cell of FliK/FlgE operon fusions analyzed using anti-flagellin immunostaining. The number of analyzed, individual bacteria (n) and the average number of flagella per cell ± SD is indicated. WT, wildtype; SD, standard deviation.

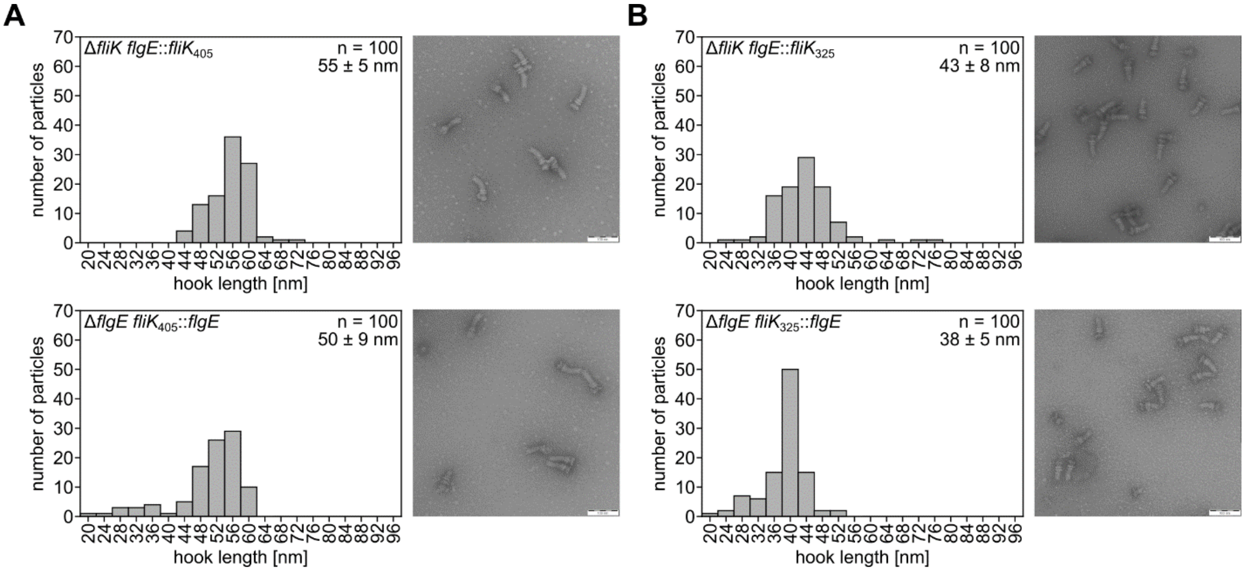


**Supplemental Figure S8. Hook-lengths of FliK/FlgE operon fusions.** Hook-length distribution of (A) FliK$_{405}$/FlgE and (B) FliK$_{325}$/FlgE operon fusions. Left panels: Hook-length distribution of purified HBBs of FliK/FlgE operon fusions. The number of measured HBBs (n) and average hook-length ± SD is indicated. Right panels: Representative electron micrographs of purified HBBs. Scale bar = 100 nm. WT, wildtype; HBB, hook-basal body; SD, standard deviation.
